# Intestinal parasitic infection among household contacts of primary cases, a comparative cross-sectional study

**DOI:** 10.1101/723494

**Authors:** Berhanu Elfu Feleke, Melkamu Bedimo Beyene, Teferi Elfu Feleke, Tadesse Hailu Jember, Bayeh Abera

**Affiliations:** Department of Epidemiology and Biostatistics, University of Bahir Dar, Bahir Dar, Ethiopia; Department of pediatrics, St Paul University, Addis Ababa, Ethiopia; Department of Medical Laboratory Science, College of Medicine and Health Sciences, Bahir Dar University, Bahir Dar City, Ethiopia; Department of Microbiology, College of Medicine and Health Sciences, Bahir Dar University, Ethiopia

**Keywords:** intestinal parasite, contact screening, secondary attack rate, household members

## Abstract

**Background:** Intestinal parasitic infection affects 3.5 billion people in the world and mostly affecting the low socio-economic groups. The objectives of this research were to estimate the prevalence and determinants of intestinal parasitic infection among family members of known intestinal parasite infected patients.

**Methods and materials:** A comparative cross-sectional study design was implemented in the urban and rural settings of mecha district. The data were collected from August 2017 to March 2019 from intestinal parasitic infected patient household members. Epi-info software was used to calculate the sample size, 4531 household members were estimated to be included. Data were collected using interview technique and colleting stool samples from each household contact of intestinal parasite patients. Descriptive statistics were used to estimate the prevalence of intestinal parasites among known contacts of intestinal parasites patients/family members. Binary logistic regression was used to identify the determinant factors of intestinal parasitic infection among family members.

**Results:** The prevalence of intestinal parasite among household contacts of parasite-infected family members was 86.14 % [95% CI: 86.14 % - 87.15 %]. *Hookworm* parasitic infection was the predominant type of infection (18.8%). Intestinal parasitic infection was associated with sex, environmental sanitation, source of water, habit of playing with domestic animals, the presence of chicken in the house, the presence of household water filtering materials, overcrowding, personal hygiene, residence, and substandard house, role in the household, source of light for the house, floor materials, trimmed fingernails, family size, regular hand washing practice, barefoot.

**Conclusion:** The prevalence of intestinal parasites was high among household contacts of primary confirmed cases.

## Introduction

Intestinal parasites are groups of worm’s primary affecting the gastrointestinal tracts broadly contains flatworms (tapeworms and flukes) and roundworms (ascariasis, pinworm, and hookworm infections)[1]. The mode of transmission includes ingestion of uncooked animal products, consuming infected water, absorption through the skin and fecal-oral [2]. Predominantly intestinal parasitic infection transmitted through feco-oral route [3]. That means all family members living in intestinal positive patients at higher risk of acquiring the infection.

A patient infected with intestinal parasite manifests with abdominal cramp, vomiting, excessive bowl sound, nausea, diarrhea, loss of appetite, malabsorption, skin itching [4]. Due to unspecified symptoms, the diagnosis of intestinal parasitic infection usually performed by taking stool samples and applying different laboratory techniques, concentration technique is more valid than the other laboratory techniques [5].

Intestinal parasitic infection affects 3.5 billion people in the world and mostly affecting the low socio-economic groups [6]. Soil-transmitted helminths infection (Ascaris lumbricoid, Trichuris trichiura and hookworm) alone affects 1.5 billion people worldwide [7]. Sub-Saharan Africa bears the highest burden for both helminths infection and other intestinal parasitic infections [8].

The complications of intestinal parasites include malnutrition, intestinal obstruction, growth retardation, immunodeficiency and affecting the socioeconomic development of the nations [9]. Intestinal parasitic infection was associated with gender, age and role in the household, socioeconomic characteristics, levels of education, poor sanitation, proximity to water sources, family size, environmental sanitation, hand washing practice, untrimmed fingernail, housing conditions, resident, barefoot [10-18]

The management of intestinal parasitic infection was not complicated and most intestinal parasitic infection can be effectively treated with a single dose anti-helminths. However, the intestinal parasitic intervention neglects the household contacts because there is no available evidence on the prevalence of intestinal parasites among household members; so, this study was conducted to give baseline evidence on the estimate of household secondary cases.

The objective of this research work was to estimate the prevalence and determinants of intestinal parasitic infection among family members of known intestinal parasitic infected patients.

## Methods and materials

The comparative cross-sectional study design was implemented in the urban and rural settings of mecha district. Mecha district was located in the north-west of Ethiopia and the district contains 10 health centers and 1 general hospital. The data were collected from August 2017 to March 2019. Data were collected from intestinal parasitic infected patient household members.

The sample size was calculated using Epi-info software version 7 using the assumption of 95 % CI, power of 85, rural to an urban ratio of 2, none response rate of 10% gives 1510 household members from the urban setting and 3021 household members from the rural settings.

Household members were selected using contact tracing. Patient diagnosed positive for parasitic infection in the district health facility were used to trace for their family members intestinal parasitic infection status. All family members were screened for intestinal parasitic infection. Data were collected using interview technique and collecting stool samples from each household contact of intestinal parasite patients. Clinical nurses were recruited for the data collection phase during interview and health officers were recruited for supervision. The stool samples were collected from each family member of known intestinal parasitic infected patients and transported to the nearby health facility for the analysis. From each known contact, one gram stool sample was collected in 10 ml SAF (sodium acetate-acetic acid-formalin solution). Formal ether concentration technique was used to identify the presence of intestinal parasites. The stool sample was well mixed and filtered using a funnel with gauze. Around 7 ML (Milliliter) normal saline and 3 ml of ether were added, mixed well and then centrifuged for 5 minutes at 2000 RPM. Finally, the supernatant was discarded and the sediment was examined for parasites under the microscope [19].

Data were entered to Epi-info software and transported to SPSS for analysis. Descriptive statistics were used to estimate the prevalence of intestinal parasites among known contacts of intestinal parasites patients/family members. Binary logistic regression was used to identify the determinant factors of intestinal parasitic infection among family members. Hand washing practice was measured if the participants wash his/her hands after visiting the toilet, before cooking food and before feeding.

Ethical clearance was obtained from research and ethical review board from (institutional research review board) collage of medicine and health sciences, Bahir Dar University. Permission letter was obtained from Amhara National Regional State Health Bureau ethical committee and Mecha district health office. Written informed consent was obtained from each study participants or guardians. Those study participants with intestinal parasites were referred to the nearby health facility for further management. The confidentiality of the data was kept at all stages.

## Results

A total of 4436 study participants were included giving for the response rate of 98 %. Female constitute 50% of the study participants, and 67% of the study participants were from the rural area. (Table 1)

**Table 1:** Population profile of the study participants (n=4436)

| SN<sup>1</sup> | Population profile |  | Frequency | Percentage |
| --- | --- | --- | --- | --- |
| 1. | Sex | Female | 2206 | 49.7 |
|  |  | Male | 2230 | 50.3 |
| 2. | Environmental sanitation | Clean | 1323 | 29.8 |
|  |  | Dirty | 3113 | 70.2 |
| 3. | Source of light for the house | Modern | 1073 | 24.2 |
|  |  | Traditional | 3363 | 75.8 |
| 4. | Floor materials of the house | Mud | 3190 | 71.8 |
|  |  | Others | 1246 | 28.2 |
| 5. | Household water filtering mechanisms | Present | 861 | 19.4 |
|  |  | Absent | 3575 | 80.6 |

<sup>1</sup> Serial number

|  |  |  |  |  |
| --- | --- | --- | --- | --- |
| 6. | Fingernails of the respondents | Trimmed | 927 | 20.9 |
|  |  | Not trimmed | 3509 | 79.1 |
| 7. | Family size | $\leq 4$ | 661 | 14.9 |
| | | $> 4$ | 3775 | 85.1 |
| 8. | Educational status | Illiterate | 1744 | 39.3 |
|  |  | Formal education | 2557 | 57.6 |
|  |  | Informal education | 135 | 3 |
| 9. | Resident | Rural | 2960 | 66.7 |
|  |  | Urban | 1476 | 33.3 |
| 10. | Marital status | Single | 3320 | 74.8 |
|  |  | Married | 1056 | 23.8 |
|  |  | Divorced | 42 | 0.9 |
|  |  | Widowed | 18 | 0.4 |
| 11. | Age in years | 0-10 | 1744 | 39.3 |
|  |  | 11-20 | 2035 | 45.9 |
|  |  | 21-30 | 215 | 4.8 |
|  |  | 31-40 | 303 | 6.8 |
|  |  | 41-50 | 12 | 0.3 |
| | | $> 50$ | 127 | 2.9 |

The prevalence of intestinal parasitic infection among family members was 86.14 % [95% CI: 86.14 % - 87.15 %]. Hookworm parasitic infection (18.8%) was the predominant parasitic infection followed by *Enatmeba histolytic* (11.4%), 36.2 % of family member has a heavy intensity of infection (Table 2).

**Table 2:** The type of parasitic infection among household members (n=4436).

| Intestinal parasitic species | Frequency | Percent |
| --- | --- | --- |
| Not infected | 615 | 13.9 |
| *Hookworm* | 834 | 18.8 |
| *Ascaris lumbricoid* | 375 | 8.5 |
| *S. mansoni* | 198 | 4.5 |
| *Trichuris Trichiura* | 332 | 7.5 |
| *E. histolytica* | 505 | 11.4 |
| *Balantidium Coli* | 411 | 9.3 |
| *G. lamblia* | 302 | 6.8 |
| *Hymenolepis nana* | 29 | .7 |
| Mixed infections | 835 | 18.8 |

### Intestinal Parasitic infection among children

The prevalence of intestinal parasitic infection among children family members was 82.77 % [95% CI: 81.08 % −84.47 %]. After adjusting for sex, environmental sanitation, source of light for the house, floor material, the presence of water filtering materials, size of the fingernails, barefoot, family size, source of water, overcrowding, personal hygiene, the presence of chicken in the house, and substandard house: Intestinal parasitic infection among household members was associated with sex, environmental sanitation, source of water, habit of playing with domestic animals, the presence of chicken in the house, the presence of household water filtering materials, overcrowding, personal hygiene, residence, and substandard house (Table 3)

**Table 3:** The determinants of intestinal parasitic infection among children household members (n=1904).

| Variable |  | IP |  | COR [ 95 % CI] | AOR [ 95 % CI] | p-value |
| --- | --- | --- | --- | --- | --- | --- |
|  |  | Infected | Not infected |  |  |  |
| Sex | Male | 717 | 168 | 0.79 [0.62-1.02] | 0.76[0.58-0.99] | 0.04 |
|  | Female | 859 | 160 |  |  |  |
| Environmental sanitation | Clean | 168 | 10 | 3.79 [1.92-7.71] | 0.04 [0.01-0.14] | <0.01 |
|  | Dirty | 1408 | 318 |  |  |  |
| Household water filter | Present | 601 | 105 | 1.31 [1.01-1.70] | 0.28 [0.18-0.44] | <0.01 |
|  | Absent | 975 | 223 |  |  |  |
| Habit of playing with domestic animals | Present | 1166 | 261 | 0.73 [0.54-0.99] | 1.62 [1.08-2.45] | 0.02 |
|  | Absent | 410 | 67 |  |  |  |
| Chicken in the household | Present | 1069 | 256 | 0.59 [0.44-0.79] | 4.42 [2.81-6.95] | <0.01 |
|  | Absent | 507 | 72 |  |  |  |
| Water source | Pipe | 443 | 234 | 0.16 [0.12-0.21] | 0.05 [0.03-0.07] | 0.03 |
|  | Others | 1133 | 94 |  |  |  |
| **Overcrowding** | Present | 956 | 152 | 1.79 [1.40-2.28] | 2.14 [1.6-2.88] | 0.01 |
|  | Absent | 620 | 176 |  |  |  |
|  | Clean | 1395 | 312 | 0.4 [0.22-0.68] | 0.26 [0.07-0.93] | 0.04 |
| **Personal hygiene** | Not clean | 181 | 16 |  |  |  |
| **Resident** | Urban | 576 | 92 | 1.48 [1.13-1.94] | 2.68 [1.86-3.89] | <0.01 |
|  | Rural | 1000 | 236 |  |  |  |
|  | Yes | 237 | 42 | 1.21 [0.84-1.74] | 1.92 [1.03-3.6] | 0.04 |
| **Substandard house** | no | 1339 | 286 |  |  |  |

### Intestinal parasitic infection in adult household members

The prevalence of intestinal parasitic infection among household members whose age greater than 16 years was 88.67% [95% CI: 87.43 % −89.90%]. After adjusting for sex, role in the household, environmental sanitation, source of light for the house, floor materials of the house, habit of ingesting raw vegetables, the presence of household water filtering materials, trimmed fingernails, substandard house, habit of playing with domestic animals, family size, the presence of chicken in the house, handwashing behavior, source of water, overcrowding, barefoot, personal hygiene, residence and chronic illness: intestinal parasitic infection among household members was associated with sex, role in the household, environmental sanitation, source of light for the house, floor materials, the presence of household water filter, trimmed fingernails, substandard house, habit of playing with domestic animals, family size, the presence of chicken in the house, regular hand washing practice, source of water for the house, barefoot, personal hygiene, resident (Table 4).

**Table 4:** The determinants of intestinal parasitic infection among adults household members (n=2532).

| Variable |  | IP |  | COR [ 95 % CI] | AOR [ 95 % CI] | p-value |
| --- | --- | --- | --- | --- | --- | --- |
|  |  | Positive | Negative |  |  |  |
| Sex | Male | 1079 | 266 | 0.07 [0.05-0.12] | 0.04 [0.02-0.09] | <0.01 |
|  | Female | 1166 | 21 |  |  |  |
| Environmental sanitation | Clean | 1280 | 107 | 2.23 [1.72-2.90] | 0.18 [0.12-0.27] | 0.01 |
|  | Dirty | 965 | 180 |  |  |  |
| Water filter | Present | 74 | 81 | 0.09 [0.06 - 0.12] | 0.21[0.12-0.4] | <0.01 |
|  | Absent | 2171 | 206 |  |  |  |
| Habit of playing with domestic animals | Present | 1670 | 149 | 2.69 [2.08 - 3.48] | 4.39 [2.58-7.47] | <0.01 |
|  | Absent | 575 | 138 |  |  |  |
| Chicken | Present | 1454 | 63 | 6.54 [4.83 – 8.85] | 3.59 [2.38-5.41] | <0.01 |
|  | Absent | 791 | 224 |  |  |  |
| Water source | Pipe | 1499 | 119 | 2.84 [2.19 – 3.67] | 0.16 [0.1-0.29] | <0.01 |
|  | Others | 746 | 168 |  |  |  |
| **Role in the household** | Children or mothers | 1277 | 39 | 8.39 [5.85-12.07] | 2.75 [1.51-4.99] | 0.01 |
|  | Others | 968 | 248 |  |  |  |
|  | Clean | 2113 | 270 | 1.01 [0.58 - 1.74] | 0.04 [0.01-0.12] | <0.01 |
| **Personal hygiene** | Not clean | 132 | 17 |  |  |  |
| **Resident** | Urban | 719 | 89 | 1.05 [0.8-1.38] | 2.32 [1.5-3.55] | <0.01 |
|  | Rural | 1526 | 198 |  |  |  |
|  | Yes | 946 | 108 | 1.21 [0.93-1.57] | 4.09[2.44-6.87] | <0.01 |
| **Substandard house** | no | 1299 | 179 |  |  |  |
| Source of light for the house | Traditional | 1692 | 247 | 0.5 [0.34-0.71] | 2.28 [1.19-4.37] | <0.01 |
|  | Modern | 553 | 40 |  |  |  |
| Family size | >4 | 1946 | 158 | 5.31 [4.05-6.97] | 7.18 [3.89-13.37] | <0.01 |
|  | ≤4 | 299 | 129 |  |  |  |
| Regular hand washing practice | Present | 208 | 2037 | 0.6 [0.41-0.87] | 0.4 [0.2-0.79] | <0.01 |
|  | Absent | 42 | 245 |  |  |  |
| Barefoot | Yes | 1499 | 119 | 2.84 [2.19-3.67] | 4.5 [2.9-6.8] | <0.01 |
|  | No | 746 | 168 |  |  |  |

## Discussion

The prevalence of intestinal parasitic infection among family members was 86.14 % [95% CI: 86.14 % - 87.15 %]. The prevalence of intestinal parasitic infection among in children family members was 82.77 % [95% CI: 81.08 % −84.47 %]. The prevalence of intestinal parasitic infection among household members whose age greater than 16 years was 88.67% [95% CI: 87.43 % −89.90%]. This finding was higher as compared to finding from England [20]. This might be due to the difference in the living condition. Our study area contains numerous contacts which increase the risk of acquiring intestinal parasites infection.

The odds of intestinal parasitic infections among female household members were 24% higher during childhood and 96% higher during adulthood. This finding agrees with other scholars works [21]. This is due to the fact that women in the household are responsible to care for the child and dispose of the waste of the child which increases their risk of acquiring the infection easily [22].

Environmental sanitation decreases the odds of intestinal parasitic infection by 96% during childhood and by 82% during adulthood. This finding agrees with finding from other parts of Ethiopia [23]. This is because environmental sanitation illuminates the reservoir for intestinal parasitic infection which finally blocks the infectious cycle of the parasites [24].

Household water filtering materials decrease the odds of intestinal parasitic infection by 72% in children and 79% in adults. This finding agrees with finding from systematic review pools across the globe [25]. This is because water treatment at the households levels eliminates the eggs/cysts of intestinal parasites from the water[26].

A habit of playing with domestic animals increases the odds of intestinal parasitic infection by 4.39 folds higher in children and 1.62 folds in adults. This finding agrees with finding from Canada [27]. This is because most intestinal parasitic infections are zoonotic in nature [28].

The presence of chicken in the household increases the odds of intestinal parasitic infection by 4.42 folds higher in children and by 3.39 folds higher in adults. This finding agrees with findings from China[29]. This is because chickens act as a reservoir to numerous species of intestinal parasites [30].

Using pipe water decreases the odds of intestinal parasitic infection by 95% in children and by 84 % in adults. This finding agrees with finding from Brazil [31]. This indicated that untreated water is a potential source of intestinal parasites infection [32].

The odds of intestinal parasitic infection were 2.75 higher in children and mothers as compared to other household members. This finding agrees with findings from Accra[33]. This is because of the proximity of mothers and children to the household wastes which harbors numerous intestinal parasites [34].

The odds of intestinal parasitic infection were 2.68 folds higher among urban children and 2.32 folds higher in the urban adults. This finding agrees with findings from India [35]. This might be due to poor environmental sanitation with the overcrowding situation in urban area [36].

Personal hygiene decreases the odds of intestinal parasitic infection by 74 % lower in children and 96 % lower in adults. This finding agrees with systematic review report from the globe [37]. This is because personal hygiene breaks the chain of intestinal parasitic infection [38].

Substandard housing increases the odds of intestinal parasitic infection by 1.92 folds higher in children and by 4 folds higher in adults. This finding agrees with finding from Brazil [39]. This is because people living under a better housing condition which has better sanitation facility [40].

The odds of intestinal parasitic infection were 2.28folds higher among household members using traditional light for their house. This finding agrees with clinical trial results [41]. This is because if the household was supplied with electricity, the household members can become aware of a health-related condition thought radio, television mass education which finally increases their awareness of a health related condition.

Regular hand washing practice decreases the odds of intestinal parasitic infection by 60 % lower. This finding was in line with 2018 finding from Ethiopia [42]. This is because regular hand washing practice breaks the life cycles of intestinal parasitic infection from an infected host to susceptible host[43].

Higher family size increases the odds of intestinal parasitic infection by 7.18 folds higher. This finding agrees with the previous finding from the same study area[44]. This is because high family size decreases the access to the basic sanitary facility due to sharing of the limited resources.

Barefoot increases the odds of intestinal parasitic infection by 4.5 folds higher. This finding was in line with 2018 results from Nigeria [45]. This is because barefoot allows the entry of intestinal parasites like hookworm at its infective stage [46].

The main limitation of this study was a failure to identify the incident and prevalent cases, but the overall aim of this study was to estimate the prevalence of intestinal parasitic infection among household members mixing of new or pre-existing cases will not create a huge problem.

## Conclusion

The burden of intestinal parasites was high among household contacts of intestinal parasite infected family members. Intestinal parasitic infection among household members was determined by gender, environmental sanitation, household water treatment, habit of playing with domestic animals, The presence of chicken in the household, source of water, role in the household, resident, housing condition, source of light for the house, hand washing practice, family size, and barefoot.

## Recommendation

Clinicians must trace and care for all household contacts of intestinal parasite patients in order to make the interventions effective.

## Acknowledgments

Our heartfelt acknowledgment goes to household members for good cooperation during the field work. We would also like to acknowledge Mecha district health office for their unreserved efforts. At last but not least we would also like to acknowledge all organization and individuals that contributed to this research work.

